# Linearly polarized excitation enhances signals from fluorescent voltage indicators

**DOI:** 10.1101/2021.07.21.453006

**Authors:** William Bloxham, Daan Brinks, Simon Kheifets, Adam E. Cohen

**Affiliations:** Department of Chemistry and Chemical Biology, Harvard University, Cambridge, MA 02138, USA; Department of Physics, Harvard University, Cambridge, MA 02138, USA; Physics of Living Systems, Department of Physics, Massachusetts Institute of Technology, Cambridge, MA 02139, USA; Department of Imaging Physics, Delft University of Technology, Delft, The Netherlands; Department of Molecular Genetics, Erasmus University Medical Center, Rotterdam, The Netherlands

## Abstract

Voltage imaging in cells requires high-speed recording of small fluorescent signals, often leading to low signal-to-noise ratios. Because voltage indicators are membrane-bound, their orientations are partially constrained by the plane of the membrane. We explored whether tuning the linear polarization of excitation light could enhance voltage indicator fluorescence. We tested a panel of dye and protein-based voltage indicators in mammalian cells. The dye BeRST1 showed a 73% increase in brightness between the least and most favorable polarizations. The protein-based reporter ASAP1 showed a 22% change in brightness, and QuasAr3 showed a 14% change in brightness. In very thin neurites expressing QuasAr3, improvements were anomalously large, with a 170% increase in brightness between polarization parallel vs perpendicular to the dendrite. Signal-to-noise ratios of optically recorded action potentials were increased by up to 50% in neurites expressing QuasAr3. These results demonstrate that polarization control can be a facile means to enhance signals from fluorescent voltage indicators, particularly in thin neurites or in high-background environments.

## Introduction

Electrical signaling is the central language of the nervous system, but historically membrane voltage has been difficult to measure. Optical measurements of membrane voltage are emerging as a powerful tool for mapping bioelectric effects in neurons,^1–5^ cardiomyocytes^6,7^, and other electrically active cell types.^8,9^ Voltage indicators have been designed around protein^10,11^, organic dye^12–14^, and hybrid protein/dye^3,15^ scaffolds. Development of improved voltage indicators is a subject of much ongoing research.^16,17^

Despite recent advances, in many cases the signal-to-noise ratio (SNR) for voltage measurements remains low. The challenges are that the signals are brief (~1 ms for a typical action potential), some reporters are very dim (e.g. those based on direct retinal fluorescence have ~1% quantum yield), and fractional changes in fluorescence are often small (~50% for an action potential for the most sensitive reporters). These challenges are compounded by the tissue environment, in which voltage-dependent signals are scattered by tissue and compete with background fluorescence. Methods for increasing the SNR would expand the capabilities of voltage imaging, potentially allowing for higher temporal resolution, recording from subcellular domains (e.g. dendrites), or recording deeper in tissue.

Improvements in signal quality could come equally from improved molecular reporters or from improved optical instrumentation. Compared to the efforts on the molecular reporters, less effort has gone into developing optical systems optimized for voltage imaging in tissue. The physical attributes of voltage indicators suggest that the optimal voltage imaging system might be substantially different from systems designed for imaging reporters of other modalities, e.g. calcium.

All voltage indicators, regardless of mechanism, are physically localized to the cell membrane, a two-dimensional manifold embedded in a three-dimensional tissue. Useful signals only come from optical excitation targeted directly to the membrane: excitation either inside or outside of the target cell contributes to sample heating, background fluorescence, and phototoxicity, but not to signal. Efforts to localize two-photon (2P)^5,18^ or one-photon (1P)^1,19,20^ illumination to the membrane led to dramatic improvements SNR.

Membrane localization also raises the possibility of orientational order in voltage-reporting chromophores. Chromophores preferentially absorb light when the polarization of the excitation is aligned with the transition dipole between the ground and electronically excited state. By optimizing this alignment at a patch of membrane, one might increase the in-focus fluorescence signal, without affecting the background (in which chromophores are assumed to be randomly oriented), and thereby improve the SNR.

The degree of improvement in fluorescence signal depends on the degree of orientational order in the membrane-bound chromophores and on the physical structure of the membranes of interest. In this paper we explore theoretically and experimentally the prospects for enhancing SNR by imaging with polarized optical excitation. We consider the organic dye indicator BeRST1^12^, and the protein-based indicators ASAP1,^11^ QuasAr3,^1^ ArcLight,^21^ Ace-mNeonGreen,^22^ and CAESR.^23^ Up to 73% enhancements in signal are observed for BeRST1 in cell bodies and 170% increases for QuasAr3 in thin neurites. These findings show that simple changes in optics could substantially enhance fluorescent voltage signals without increasing background.

## Theory

Brasselet and coworkers have presented a detailed study of the responses of chromophores in membranes to polarized 1-P excitation.^24^ Here we review the key results relevant to voltage imaging. When a fluorophore with excitation transition dipole **μ** is excited by linearly polarized light with polarization vector **e**, the fluorescence *F* is proportional to |**μ** · **e**|^2^. Modulation of optical excitation by changes in linear polarization is called linear dichroism (LD), and when the excitation is monitored via fluorescence, the phenomenon is called fluorescence detected linear dichroism (FDLD).

We assume that membrane-bound chromophores have free rotation about the surface normal, but fixed orientation relative to the membrane plane. The population then has a cone-shaped distribution of transition dipoles, with the cone axis of symmetry along the membrane normal (**Figure 1a**). For high numerical aperture (NA) objectives, the dipolar emission pattern is collected over enough angles that one can safely neglect the dependence of the collection efficiency on the orientation of **μ**.^25,26^

**Figure 1.**
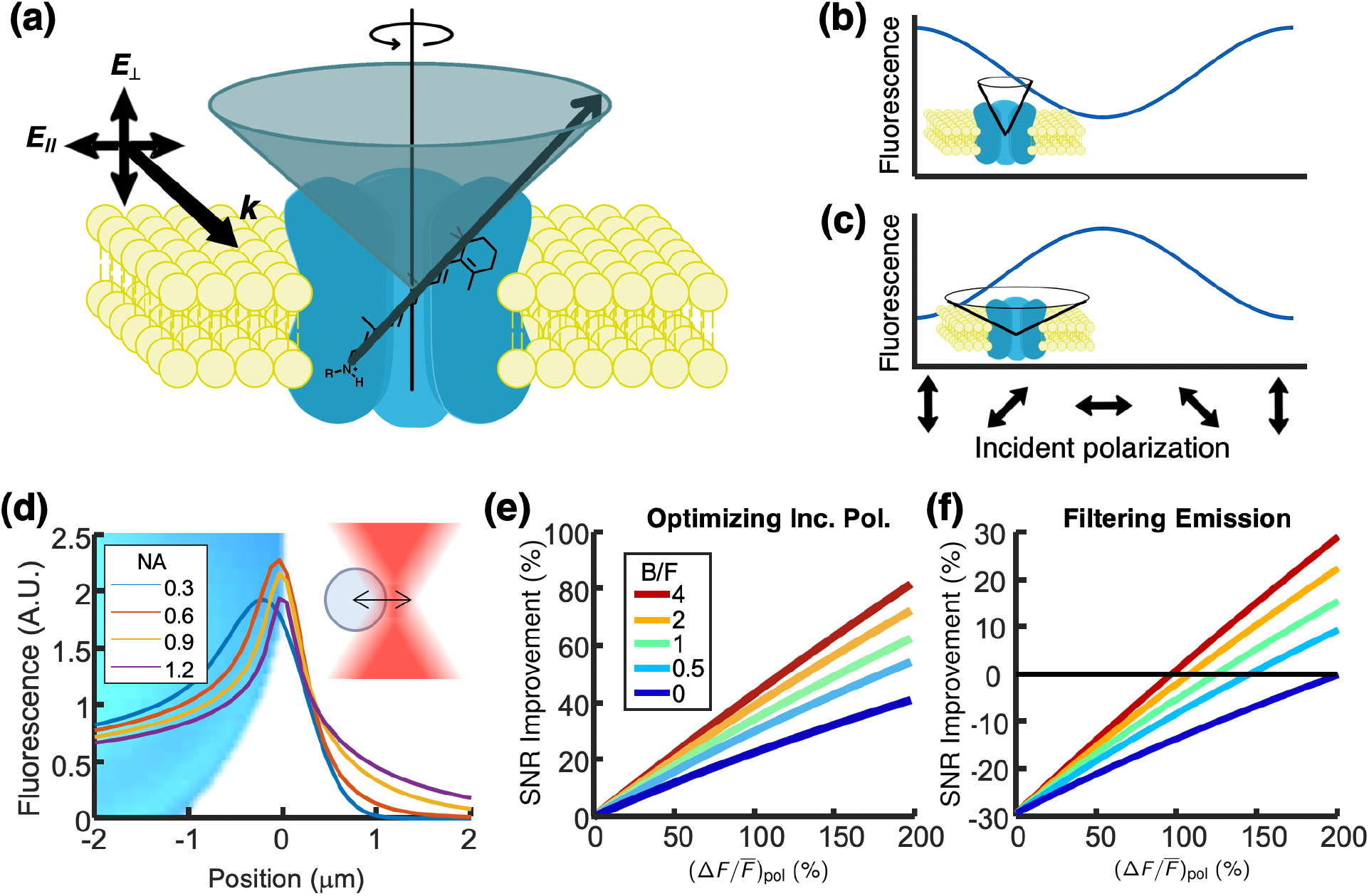
Polarized excitation can enhance signal from membrane-bound indicators. (a) Geometry of a voltage-indicating chromophore in a membrane. An ensemble of chromophores has a cone-shaped distribution of transition dipoles. (b) When light propagates in the plane of the membrane, the efficiency of fluorescence excitation depends on the polarization. Out-of-plane polarization favors excitation for chromophores with a narrow cone-angle and (c) in-plane polarization favors excitation for chromophores with a wide cone angle. (d) Total fluorescence emission from a 10 μm-diameter spherical shell illuminated with an unpolarized diffraction-limited Gaussian beam. Total fluorescence is relatively insensitive to illumination numerical aperture (NA), but is substantially enhanced at the equator of the sphere. (e) Theoretical shot noise-limited SNR improvement from optimizing the polarization of the incident light (compared to the expectation value of the SNR under a random linear polarization). (f) Theoretical SNR improvement from filtering the emission by polarization. B/F is the background fluorescence divided by the fluorescence from the indicator. Legend applies to (e) and (f).

For illumination traveling orthogonal to the plane of the membrane, there can be no FDLD signal, due to the azimuthal symmetry of the distribution of **μ**. However, when illuminating a cell from above, the membrane around the equator is illuminated edge-on and offers the prospect of producing an FDLD signal. If the cone half-angle, θ_cone_, of the distribution of transition dipole orientations is narrow, then the fluorescence is maximized when the polarization of the excitation is perpendicular to the membrane (*F*_⊥_, **Figure 1b**). If θ_cone_ is wide, then the fluorescence is maximized when the polarization is in the plane of the membrane (*F*_∥_, **Figure 1c**).

We define the fluorescence detected linear dichroism by

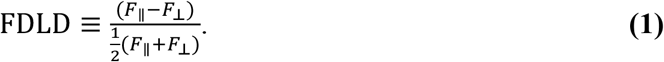

Assuming that the light is propagating entirely in the plane of the membrane, an average of the FDLD signal over all azimuthal chromophore orientations yields:

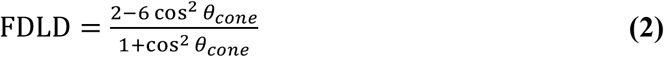

The derivation is given in the **Supporting Information**. FDLD is between −200% (when *θ_cone_* = 0°) and 200% (when *θ_cone_* = 90°). There is no FDLD signal when θ_cone_ = 54.7°, the so-called “magic angle.”^27^ An analogous calculation for two-photon excitation is given in the **Supporting Information**.

In an image of a roughly spherical cell under wide-field illumination, the fluorescence at the equatorial plane appears to be brighter than at the poles, a consequence of geometrical projection. To simulate this effect, we convolved a 10 μm diameter spherical shell (representative of a nominal cell membrane) with the point-spread functions of a series of objective lenses with different numerical apertures. **Figure 1d** shows the fluorescence enhancement at the equatorial membrane. This result demonstrates that one achieves more signal per input-photon by targeting illumination to the equatorial periphery of the cell, rather than flood-illuminating the whole cell. One should then either a) select the polarization as a function of azimuth around the cell equator to maximize the FDLD signal, or b) use illumination of a single polarization and target the excitation to the azimuthal angles where the polarization is most efficient at exciting the reporter.

The shot-noise-limited SNR in a voltage imaging experiment is:

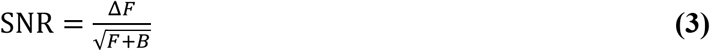

where Δ*F* is the change in fluorescence associated with the electrical event of interest, *F* is the baseline fluorescence of the reporter, and *B* is the background fluorescence. Favorable polarization alignment enhances Δ*F* and *F* proportionally, without affecting *B*. **Figure 1e** plots the predicted enhancement in SNR from favorable polarization vs. random polarization as a function of the amplitude of the *FDLD* signal, showing the possibility for nearly two-fold enhancement in SNR.

In principle, one might achieve further enhancement of signal-to-background ratio by passing the emission through an appropriately aligned polarizer too. The background fluorescence is (presumably) unpolarized, while the signal fluorescence retains some polarization, both because of selection of oriented chromophores by the polarized excitation, and by polarized emission from the inhomogeneous distribution of chromophore orientations in the membrane. A polarizer in the emission path could, in principle, increase the signal-to-background ratio by passing the polarized in-focus emission while rejecting half the unpolarized background. **Figure 1f** shows that the resulting enhancement in SNR is predicted to be small and often negative, so this approach was not considered further (calculations are in the **Supporting Information**).

## Results

### Fluorescence detected linear dichroism in HEK cells

We developed an optical system to illuminate a sample with collimated laser illumination at a variety of wavelengths, with synchronized polarization control and image acquisition (**Figure 2a**, **Methods**). We began by measuring the FDLD signals of BeRST1 (N = 2 cells), ASAP1 (N = 4 cells), QuasAr3 (N = 2 cells), ArcLight (N = 3 cells), Ace-mNeonGreen (N = 3 cells), and CAESR (N = 4 cells) in HEK cells. For each cell, we recorded wide-field epifluorescence images while alternating the polarization of the incident light between horizontal and vertical linear polarizations. We averaged the frames corresponding to each condition and calculated the FDLD signal pixel-by-pixel.

**Figure 2.**
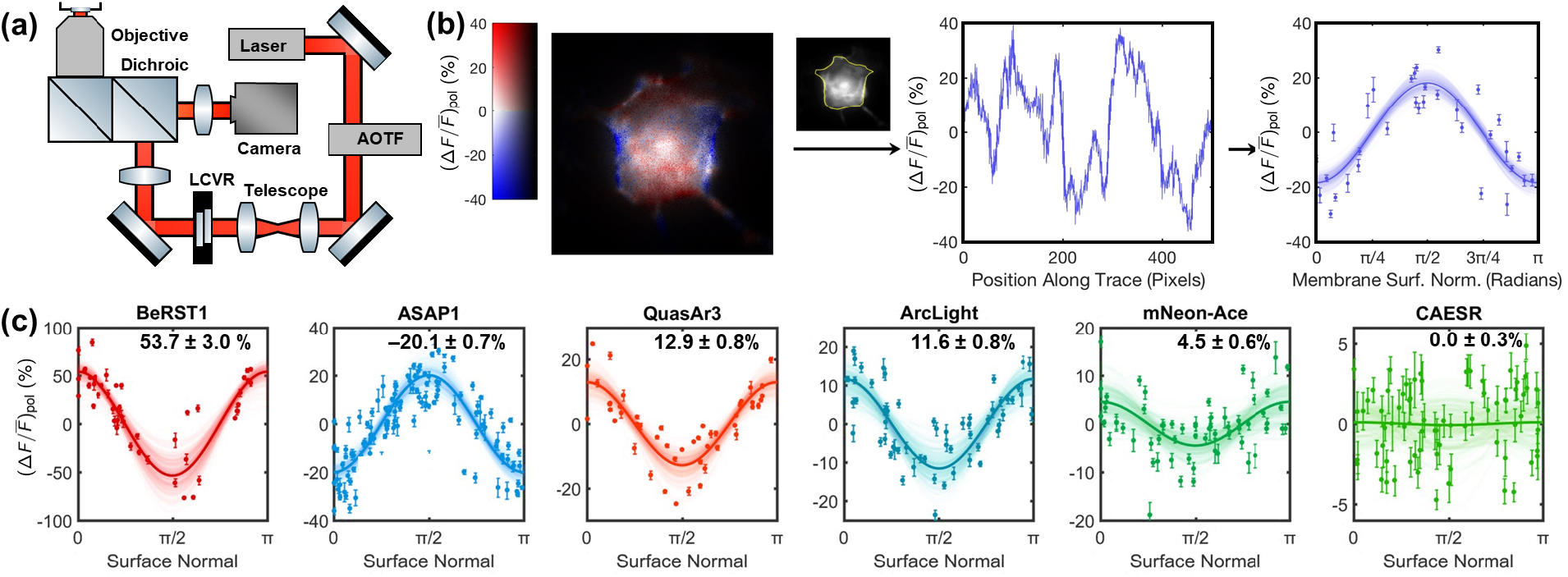
Fluorescence-detected linear dichroism in voltage-indicating chromophores. (a) Microscope used for widefield imaging experiments. Details of the setup are in Materials and Methods. The acousto-optical tunable filter (AOTF) modulated the illumination intensity and the liquid crystal variable retarder (LCVR) provided polarization control. (b) Steps for measuring linear dichroism in cells. Fluorescence images were acquired under orthogonal polarizations. The FDLD signal was extracted along the perimeter of the cell, and then mapped as a function of the direction of the membrane surface normal. (c) Linear dichroism data from HEK cells labeled with BeRST1, ASAP1, QuasAr3, ArcLight, mNeon-Ace, and CAESR. Colors correspond to approximate absorption peaks for each indicator. Each point corresponds to a line segment from the traces of the cell perimeters, and its error bar represents the standard error in the average FDLD for the pixels in that line segment.

We created a trace of the cell membranes by manually defining the vertices of a series of line segments around the periphery of each cell. We extracted the FDLD signal as a function of position along the trace and calculated the membrane surface normal for each line segment. We binned the sampled FDLD values by line segment, combined the data from all cells expressing each indicator, and fit a sinusoidal curve for FDLD as a function of membrane surface normal (**Methods** and **Supporting Information**). The process of mapping the orientation-dependent FDLD signal is illustrated for a single ASAP-expressing cell in **Figure 2b**.

**Figure 2c** shows the results for each voltage indicator. The FDLD values on each panel use the sign convention from Eq. 1. Rearranging Eq. 1 shows that the ratio of fluorescence with polarization optimal vs orthogonal is given by:

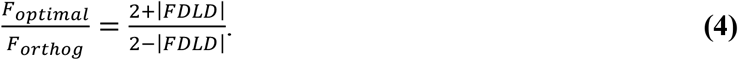

This ratio is: BeRST1 1.73 ± 0.056., ASAP1 1.22 ± 0.009, QuasAr3 1.14 ± 0.009, ArcLight 1.12 ± 0.009, mNeon-Ace 1.046 ± 0.006, CAESR 1 ± 0.003. The voltage-sensitive dye BeRST1 and the protein-based sensor ASAP1 had the FDLD signals with the largest magnitudes, though they were of opposite sign, suggesting that the BeRST1 chromophore has an orientation close to the plane of the membrane while the ASAP1 chromophore is approximately perpendicular to the membrane. Indeed, the molecular structure of BeRST1 has a ‘T’ shape, with a molecular wire penetrating the membrane and the chromophore in the membrane plane.^12^

The QuasAr3 reporter also showed a substantial FDLD signal, a consequence of the fixed orientation of the retinal chromophore relative to the protein scaffold. The two FRET-based protein reporters, Ace-mNeonGreen and CAESR, showed small FDLD signals, presumably a consequence of tumbling of the fluorescent protein which was tethered loosely to the membrane. The values of the FDLD signals corresponded to the following ensemble-averaged cone angles: BeRST1 62 ± 4°, ASAP1 52 ± 0.1, QuasAr3 56.4 ± 0.1, ArcLight 56.3 ± 0.1, mNeon-Ace 55.4 ±0.08, CAESR 54.7 ± .04, though the values for mNeon-Ace and CAESR more likely reflect orientational averaging rather than a distribution peaked around a specific cone angle.

We combined patch clamp electrophysiology and polarization-resolved fluorescence imaging to record the mean fluorescence and FDLD as a function of membrane potential. While all the reporters showed voltage-dependent changes in mean fluorescence consistent with literature reports, we did not observe voltage-dependent changes in FDLD signals in any of the indicators. These findings suggest that voltage-induced reorientations of the chromophores were below our detection sensitivity. Previous work reported a voltage-dependent 2P FDLD signal in ArcLight.^28^ Our failure to observe this effect is likely due to the lower orientational sensitivity of 1P vs. 2P linear dichroism measurements.^29^

### Axons and dendrites

Next we considered FDLD measurements in thin neurites (axons or dendrites), where voltage imaging is particularly challenging due to the small membrane surface areas. We calculated the predicted dependence of the FDLD signal on chromophore cone angle in the cylindrical geometry of a slender neurite. We assumed that the fluorescence was averaged over the circumference of the neurite, i.e. that the transverse structure of the neurite was not resolved. The optical excitation strength, |**μ** · **e**|^2^, was a function of three angles: the cone half angle *θ_cone_* for the distribution of chromophore orientations relative to the surface normal, the azimuthal angle *ϕ*_1_ about the surface normal, and an azimuthal angle *ϕ*_2_ about the axis of the neurite. This model and these angles are illustrated in **Figure 3a**.

**Figure 3.**
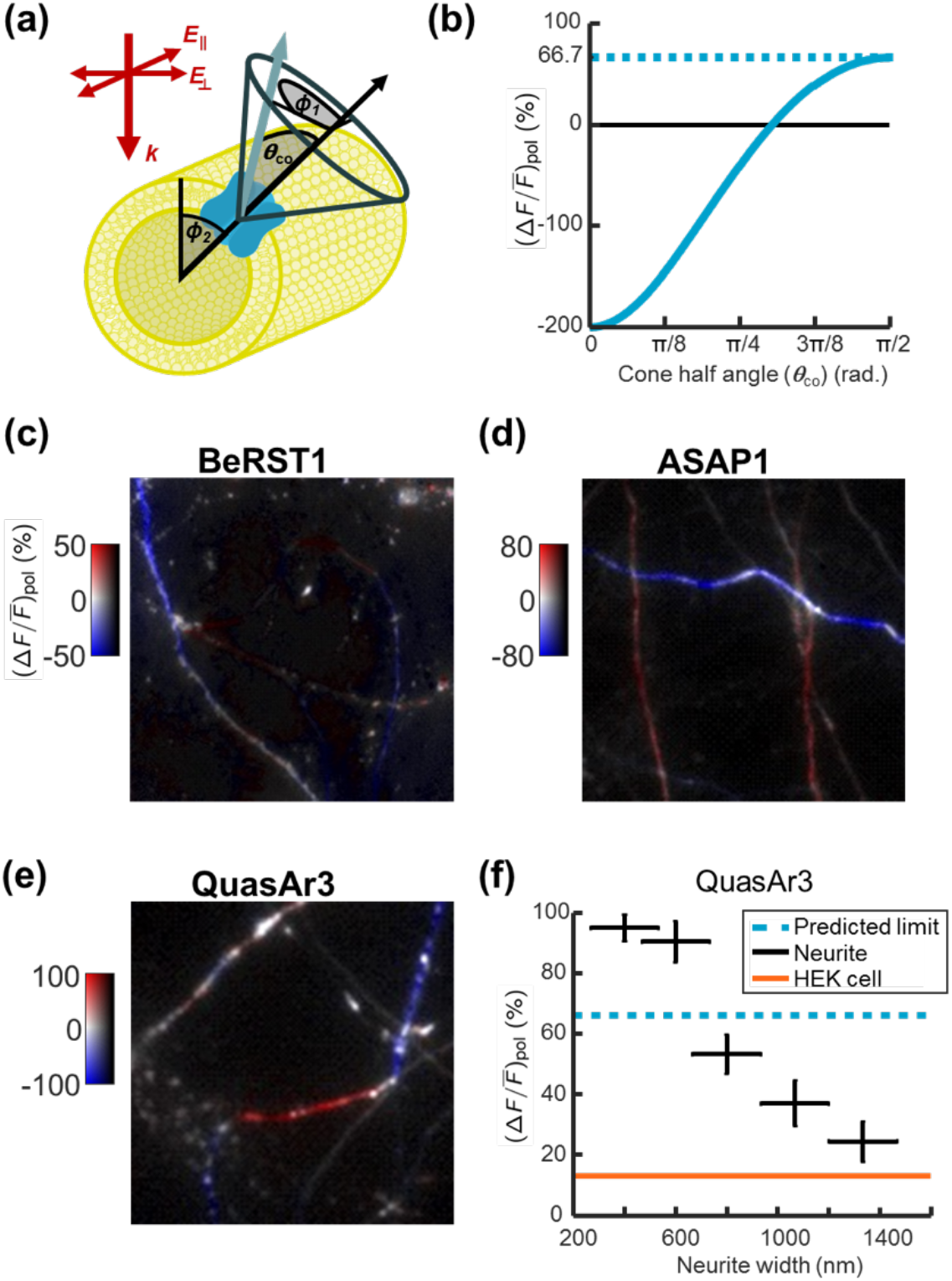
Polarized excitation enhances voltage indicator fluorescence in neurites. (a) Geometry of a neurite showing the distribution of chromophore orientations. FDLD signals are calculated by averaging over *ϕ*_1_ and *ϕ*_2_. (b) FDLD signal as a function of chromophore cone angle, assuming that photons are binned across the diameter of the neurite. FDLD images were acquired for neurons labeled with (c) BeRST1, (d) ASAP1, and (e) QuasAr3. (f) In neurites expressing QuasAr3, the FDLD signal was enhanced in thinner neurites relative to thicker ones. The orange line represents the FDLD signal measured in cell bodies. The blue dashed line represents the theoretical maximum FDLD signal, assuming isotropic in-plane chromophore orientations. Neurites were manually sorted by width into five bins (*N* = 41 neurites). The smallest bin corresponded to neurites with widths below the resolving power of the microscope. The remaining bins corresponded to neurites with widths of approximately 2, 3, 4, and 5 pixels in the recorded images. Error bars represent the width of the bins and the standard error of the mean FDLD for the neurites in each bin.

The relationship between the *θ_cone_* and the polarization anisotropy is:

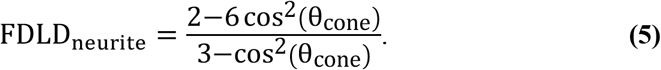

The derivation is given in the **Supporting Information**. Eq. 5 predicts that FDLD has a lower limit of −200% in the case of *θ_cone_* = 0° and an upper limit of 66.7% in the case of *θ_cone_* = 90°. This result differs from the calculations for a flat section of membrane where the bounds were – 200% and 200%. This difference arises due to contributions in the neurites from the top and bottom surfaces. A surface perpendicular to the direction of light propagation has zero FDLD, but for *θ*_cone_ > 0° the surface nonetheless contributes fluorescence. Mixing the fluorescence from the top and bottom surfaces with the fluorescence from the edges of the neurite leads to a decrease in the maximum FDLD signal when *θ*_cone_ approaches 90 degrees.

We measured linear dichroism in neurites of cultured neurons using the three indicators that exhibited the greatest FDLD signals: BeRST1, ASAP1, and QuasAr3. **Figures 3c-e** give example fields of view for each indicator. We collected 9 to 27 fields of view for each indicator (most of which contained multiple neurites and many of which contained somas in addition to neurites). We looked only at nearly vertical and nearly horizontal neurites and sorted our observations by neurite width. In the thinnest neurites (i.e. those with diameters near or below the diffraction limit), we observed FDLD values: BeRST1 30.9 ± 0.7% (N=14), ASAP1 –59.1 ± 0.4% (N=21), and QuasAr3 92.5 ± 1.2% (N=19). These FDLD values correspond to fluorescence ratios F_optimal_/F_orthog_: BeRST1 1.37 ± 0.01, ASAP1 1.84 ± 0.008, and QuasAr3 2.72 ± 0.04, i.e. there was a 172% enhancement in the QuasAr3 signal by optimizing the polarization.

The anticipated FDLD values in the neurites can be obtained by reference to Fig. 3b and the cone angles extracted from the measurements on the HEK cells. Based on the HEK cell measurements, one would anticipate FDLD values in neurites to be: 24 ± 12% for BeRST1, −10.5 ± 0.4 % for ASAP1, and 6.0 ± 0.4 % for QuasAr3. For QuasAr3 and ASAP1 the measured values are much larger than the predictions, whereas for BeRST1 the predicted and measured values were similar. In the somas of the neurons we observed FDLD values similar to those from HEK cells, indicating that the anomalously large FDLD values in neurites were not due to a neuron-specific change in *θ*_cone_.

The data for QuasAr3 were particularly striking because these results showed FDLD values in the thinnest neurites (92.5 ± 1.2%) that exceeded the maximum theoretical value, 66.7%, corresponding to θ_cone_ = 90°. While the reason for this discrepancy is not known, we speculate that it may reflect an inhomogeneous distribution of azimuthal orientations (*ϕ*_1_) in the membrane. The strong curvature around the neurite axis could break the azimuthal symmetry, favoring some values of *ϕ*_1_ over others (e.g. if the molecule itself lacks cylindrical symmetry in the membrane plane). We are not aware of examples of curvature-induced orientational order in proteins, so this idea at present remains speculation. The FDLD for ASAP1 in neurites also exceeded the anticipated value based on *θ*_cone_ measured in HEK cells, but it was within the theoretically allowed range. The FDLD for the small-molecule dye BeRST1 was as expected.

### Functional recordings in neurons

To assess whether polarized illumination could improve the signal-to-noise ratio of voltage imaging, we made optical recordings from neurites of cultured neurons. Cultured neurons were transfected with QuasAr3 and subjected to simultaneous patch clamp electrophysiology and high-speed fluorescence imaging. Action potentials were induced via current injection and voltage and fluorescence responses were recorded simultaneously. We interleaved trials with vertical and horizontal polarized illumination.

**Figure 4** shows a typical example. Fluorescence signals from the neurites showed clear polarization dependence. The degree of signal enhancement varied considerably, depending upon the diameter, orientation, and smoothness of the neurite. From neurites that were well aligned with the vertical or horizontal axes, appeared to be relatively smooth, and gave a reliable fluorescent response to action potentials, the mean enhancement in signal between the optimal and orthogonal polarizations was 29% +/− 2%, with a range from 14% to 48% (*N* = 17 neurites, 6 neurons). The mean signal-to-noise enhancement was 32% +/− 2% with a range from 20% to 50%. In some instances the improvement was such that fluorescent signals were visually discernable under the optimal polarization but not the orthogonal polarization (**Supplementary Figure 1**). Considering that this enhancement only required minimal changes in the experimental setup and did not require any changes to the molecular tools or mode of gene expression, we suggest that polarized illumination could be a valuable addition to voltage imaging efforts, particularly in thin neurites.

**Figure 4.**
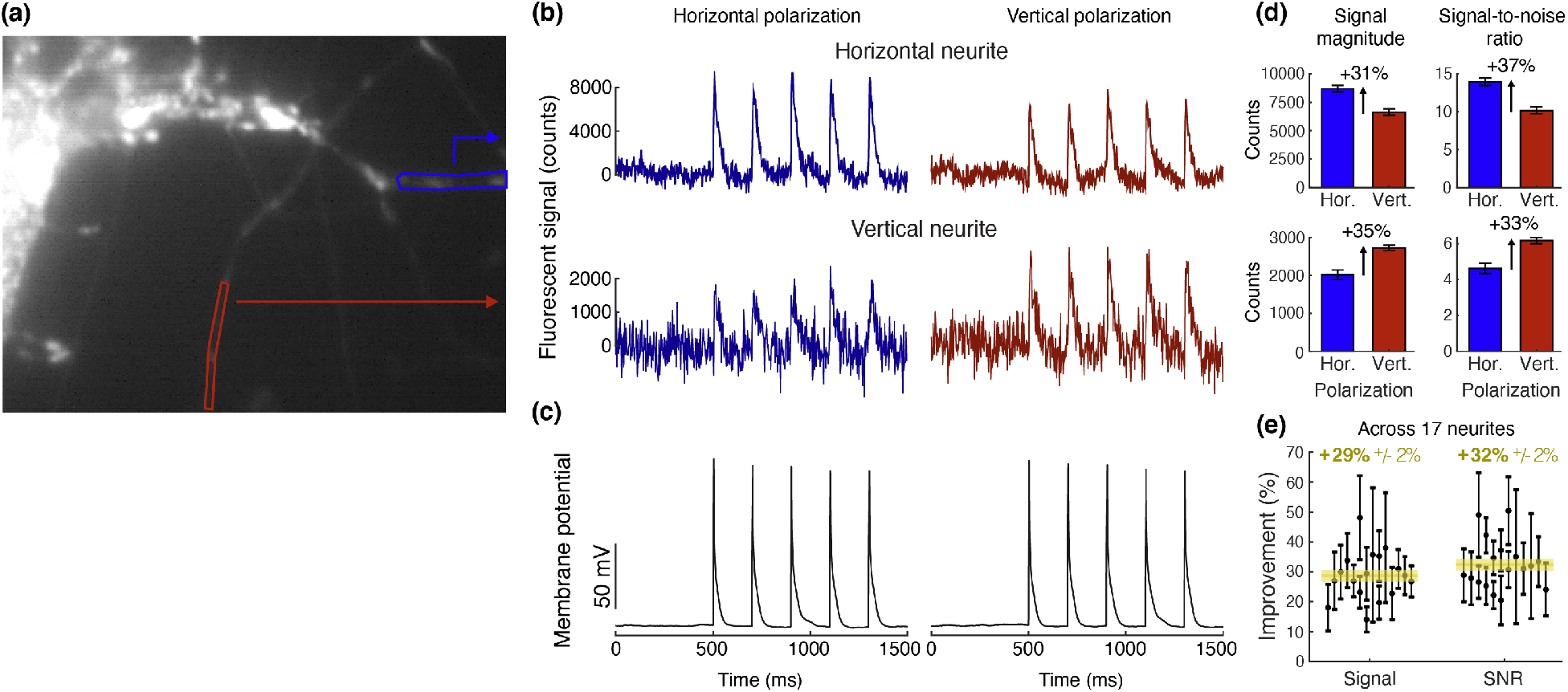
Polarized excitation enhances voltage-dependent signals in neurites. (a) Example neuron with regions indicating horizontally and vertically aligned neurites. Current injection via a patch pipette (4 ms, 400 pA, 5 Hz) triggered action potentials. (b) Fluorescence intensity traces averaged over 9 trials with each polarization. (c) Membrane potential averaged over 9 trials. (d) Signal magnitude and signal-to-noise ratios under horizontal and vertical polarizations for the neuron in (a)-(c). Background and noise were calculated as the mean and standard deviation of the fluorescence during the 500 ms before the first current pulse. The peak fluorescence in the 25 ms after the current pulse was used as the signal. Error bars represent the standard error across the five action potentials. (e) Signal magnitude and signal-to-noise ratio improvements for each of 17 neurites from 6 cells shown in black with mean improvements in yellow.

## Discussion

For most voltage imaging experiments in tissue, the background is high compared to the signal, so 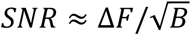. Much effort has been devoted to increasing the brightness of microbial rhodopsin-based voltage sensors. If the background *B* is due to out-of-focus reporter molecules, then increases in molecular brightness or expression level increase *F* and *B* proportionally. Eq. 3 shows that increasing the brightness by a factor *x* only increases the SNR by a factor of 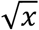. In contrast, increasing the in-focus signals *F* and Δ*F* by an amount *x* without affecting the out-of-focus background *B*, e.g. by tuning incident polarization, increases SNR by a factor of *x* in the high-background limit. Thus, in the high-background limit, a 2-fold increase in in-focus brightness is equivalent, from an SNR perspective, to a 4-fold increase in reporter brightness. These scaling arguments show that tuning incident polarization can be a potent means to enhance small voltage imaging signals.

Considerable effort is being dedicated to developing two-photon (2P) excitable voltage indicators with the hope that these tools will enable deeper imaging in intact tissue. The polarization sensitivity for 2P excitation is proportional to |**μ** · **e**|^4^, leading to a much steeper polarization dependence than for 1P excitation.^30^ Indeed polarized 2P excitation was previously used to probe the mechanism of a voltage-sensitive protein, ArcLight.^28^ In view of the strong polarization dependence, it will be important to control the polarization of the excitation, particularly when performing 2P voltage imaging in thin axons and dendrites.

Membrane-bound reporters are being developed for other modalities beside voltage, including e.g. glutamate,^31^ GABA,^32^ acetylcholine,^33^ dopamine,^34^ and serotonin.^35^ The degree of chromophore orientation in these other molecules is not known. We suggest that these other probes merit polarization analysis to determine whether signal levels can be enhanced through polarization control.

## Acknowledgments

We thank K. Williams for technical assistance with molecular biology and M. Lee for technical assistance with cell culture. This work was supported by NIH grant R01-MH117042 and the Howard Hughes Medical Institute. DB acknowledges support by an NWO Start-up Grant (740.018.018) and ERC Starting Grant (850818 – MULTIVIsion).

## Materials and Methods

### Expression of constructs in HEK293T cells

BeRST1 was a gift from Evan Miller (Berkeley); ASAP1 was a gift from Michael Lin (Stanford); Ace-mNeonGreen was a gift from Mark Schnitzer (Stanford); ArcLight was a gift from Vincent Pieribone (Yale). Addgene locations: ArcLight A242 in PCS2+, #36857; CAESR, #59172; FCK-ASAP1, #52519; QuasAr3 #107701.

HEK293T cells (ATCC; CRL-11268) were cultured and transfected as described before.^20^ Briefly, cells were grown at 37 °C, 5% CO_2_, in DMEM supplemented with 10% FBS and penicillin-streptomycin. Cells were tested negative for mycoplasma. Cells were transfected with CAESR, Quasar3 or ArcLight under the upstream CMV promoter of the FCK plasmid, and with ASAP1 under the upstream CAG promoter of the pcDNA3.1/Puro-CAG backbone. 200–400 ng of plasmid DNA was transfected using Transit 293T (Mirus) following the manufacturer’s instructions. Cells were assayed 48 hours after transfection. The day before recording, cells were re-plated onto Matrigel coated glass-bottom dishes (In Vitro Scientific) at a density of ∼10,000 cells/cm^2^.

For voltage-sensitive dye imaging, HEK cells were incubated with BeRST1 as previously described.^12^ Cells were incubated in 1 μM BeRST1 in XC buffer for 15 minutes at 37 °C and then washed to remove excess buffer.

### Neural culture

All procedures involving animals were in accordance with the US National Institutes of Health Guide for the care and use of laboratory animals and were approved by the Institutional Animal Care and Use Committee at Harvard University.

Rat glial monolayers were prepared as described previously.^36^ Briefly, 10^6^ dissociated hippocampal cells from P0 rat pups were plated on a 10-cm culture dish in glial medium, GM, composed of 15% FBS (Life), 0.4% (w/v) D-glucose, 1% GlutaMAX (Life), 1% penicillin/ streptomycin (Life) in MEM (Life). When the dish reached confluence (1–2 weeks), cells were split using trypsin onto glass-bottom dishes (In Vitro Scientific, D35-20-1.5-N) coated with poly(D-lysine) and Matrigel (BD Biosciences) at a density of 3,500 cells/cm^2^. After 3–6 days, glial monolayers were at or near confluence, and the medium was replaced by GM with 2 μM cytarabine (cytosine-β-arabinofuranoside, Sigma) to prevent further glial growth. Dishes were maintained in GM with 2 μM cytarabine until use. Dishes were discarded if microglia or neurons were identified on the monolayers.

Hippocampal neurons from P0 rat pups were dissected and cultured in neurobasal-based medium (NBActiv4, Brainbits) at a density of 30,000–40,000 neurons/cm^2^ on the pre-established glial monolayers. At 1 day in vitro (DIV), cytarabine was added to the neuronal culture medium at a final concentration of 2 μM to inhibit further glial growth.^37^ Neurons were transfected between DIV 3 and DIV 8 via the calcium phosphate transfection method.^38^ Measurements on neurons were taken between DIV 7 and 18.

### Electrophysiology

All imaging and electrophysiology were performed in extracellular (XC) buffer containing (in mM): 125 NaCl, 2.5 KCl, 3 CaCl_2_, 1 MgCl_2_, 15 HEPES, 30 glucose (pH 7.3) and adjusted to 305– 310 mOsm with sucrose. Patch clamp measurements were performed with a HEKA EPC 800 patch clamp amplifier. Filamented glass micropipettes (WPI) were pulled to a tip resistance of 5– 10 MΩ, and filled with internal solution containing (in mM):125 potassium gluconate, 8 NaCl, 0.6 MgCl_2_, 0.1 CaCl_2_, 1 EGTA, 10 HEPES, 4 Mg-ATP, 0.4 Na-GTP (pH 7.3); adjusted to 295 mOsm with sucrose. Pipettes were positioned with a Sutter MP285 manipulator. Whole-cell voltage clamp and current clamp signals were filtered at 3 kHz with the internal Bessel filter and digitized with a National Instruments PCIe 6259 board.

### Wide-Field Microscopy and Polarization Modulation

Whole-cell patch clamp and fluorescence recordings were acquired on a home-built, combined 2P and inverted epifluorescence microscope described before.^20^ The electrophysiology and optical measurements were synchronized via custom software written in LabView.

**Figure 2b** gives a schematic of the microscope pathway used for the wide-field experiments. We used 640 nm (QuasAr3, BeRST1) and 488 nm (all other indicators) Coherent Obis lasers for excitation. The laser light was passed through a Gooch and Housego AOTF and collimated and expanded using Thorlabs achromatic lenses in flip mounts for varying magnification. The light was then passed through a Liquid Crystal Variable retarder (Meadowlark LVR-200-IR1 LCVR for 640 nm excitation and Meadowlark LVR-200-VIS LCVR for 488 nm excitation) driven by a two-channel Tektronix arbitrary function generator. For 640 nm excitation, excitation light and fluorescence were separated using a Semrock FF660-Di02 dichroic mirror. For 488 nm excitation, excitation light and fluorescence were separated using a Semrock Di02-R488 dichroic mirror. Imaging was performed with an Olympus water immersion XPLN25XWMP2 objective with an NA of 1.05. Residual 640 nm laser light was rejected with a BLP01-664R-25 long-pass filter (Semrock). Residual 488 nm laser light was filtered with a BLP01-488R-25 long-pass filter. Imaging of HEK293T cells was performed on an Andor iXon X3 860 Ultra EMCCD (128×128 pixels, 24 μm pixel size); imaging of cultured neurons was performed on an Andor iXon X3 897 Ultra EMCCD (512×512 pixels, 16 μm pixel size).

### Image processing

Fluorescence signals from small neurites (*F*) were very dim, and so it was important to have an accurate calibration of the laser illumination profile and of any spurious signal sources. The two sources of spurious signal comprised background (*B*, from dark counts, ambient light) and laser-dependent signal (*L*, bleed-through and sample autofluorescence). The signal at each pixel was a sum *F* + *B* + *L*. For both HEK cell and neuron experiments, we recorded images of fluorescent cells (Fluor), calibration images without fluorescent cells (Cal), and background images with the laser light off (Back) and normalized our excitation profile as follows: Final image = (Fluor – Cal)./(Cal – Back).

### Signal processing

In the traces of Figures 2b and c small offsets were subtracted to make the FDLD curves symmetric around 0. We attribute these small asymmetries to a slight polarization dependence of the transmission of the dichroic mirrors.^39^ Error bars were calculated using the bootstrap method.

## Supplementary Calculations

### Calculations of SNR improvement from optimizing polarization

#### Optimizing excitation polarization

In this section we calculate the improvement in SNR from illuminating the sample with optimally polarized vs unpolarized light. The SNR for unpolarized illumination is

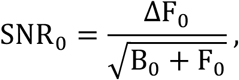

where F_0_ and B_0_ are the fluorescence and background under unpolarized illumination (here we assume that ΔF_0_ ≪ B_0_ + F_0_ , so that the change in shot noise during an event is negligible).

F_0_ is related to the fluorescence generated by the optimal (F_opt_) and orthogonal (F_orth_) linear polarizations via:

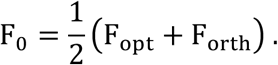

Substituting the definition of FDLD (Eq. 1), we obtain:

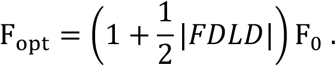

If we assume that the background is independent of polarization, then the SNR under the optimal incident polarization (OIP) is:

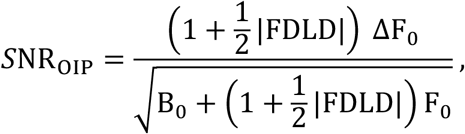

and the improvement in SNR compared to unpolarized illumination is:

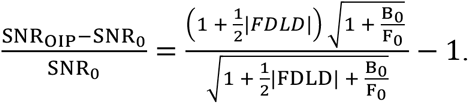

Since |*FDLD*| ≥ 0 and 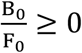, optimizing the incident linear polarization always improves the SNR.

#### Putting a polarizer in the emission path

Oriented dipoles can emit polarized light. The degree of polarization in the light that reaches the detector depends on the orientational order in the excited-state population and on the collection properties of the optical system. Here we take the degree of linear polarization at the detector as an empirical parameter and analyze the impact of inserting a polarizer in the detection path on the SNR.

In analogy to Eq. 1, we define a measure of linear dichroism in emission as:

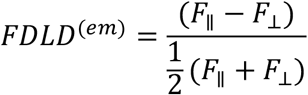

where the fluorescence values are measured with respect to the orientation of a polarizer in the emission path, and *F*_∥_ and *F*_⊥_ are the fluorescence detected with the polarizer parallel and perpendicular to the axis with greatest signal, respectively.

The total fluorescence reaching the detector is the sum of the fluorescence polarized along two orthogonal dimensions (note the factor of 2 difference from the corresponding expression for excitation):

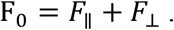

Inserting the definition of *FDLD*^(*em*)^ and rearranging yields:

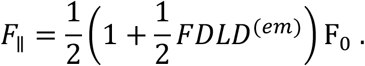

The background with filtered emission (FE) is cut in half:

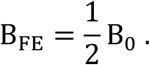

The improvement to the signal-to-noise ratio as a result of filtering the emission is

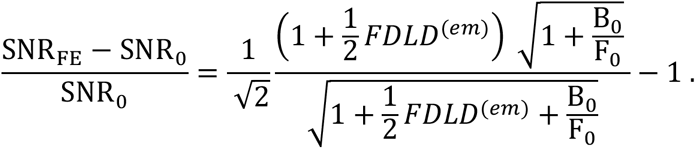

The factor 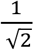 means that the improvement from filtering emission will always be less than the improvement from optimizing the incident linear polarization, and the improvement from filtering emission will not necessarily be positive. Filtering emission will be a detrimental strategy unless *FDLD*^(*em*)^ and 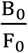 are both large.

### Calculation of FDLD for oriented chromophores in a membrane

#### Model for a flat section of membrane

Consider a planar membrane. Let the z-axis be the membrane surface normal and the y-axis be the direction of propagation of incident light. The transition dipole **μ** has (*x*, *y*, *z*) components

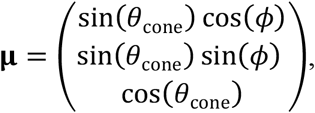

where θ_cone_ is the half-angle of the cone-shaped probability distribution for the transition dipole’s orientation and *ϕ* is the rotational position of the transition dipole within the cone-shaped position probability distribution.

The polarization vector has components

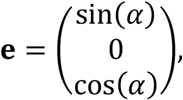

where *α* is the angle between the surface normal and the polarization of the incident light.

The fluorescence intensity as a function of θ_cone_, *ϕ*, and *α* is proportional to

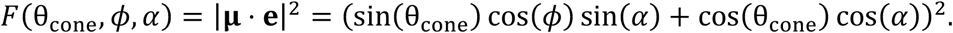

Assuming a uniform distribution for the rotational parameter *ϕ*, the expected fluorescence is proportional to

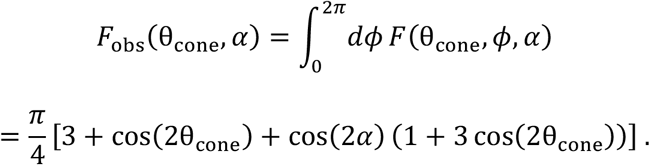

From this expression, the FDLD can be calculated as

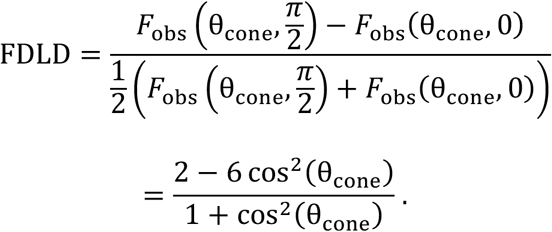

#### For 2P excitation

An analogous calculation evaluating 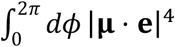 yields an FDLD signal for 2P excitation traveling in the plane of the membrane:

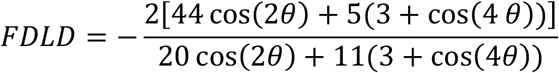

#### Model for a sub-resolution neurite

Consider a membrane tube (a “neurite”) aligned with the x-axis, illuminated by light propagating along the y-axis. We assume that all fluorescence signals are averaged across the width of the neurite. The components of the transition dipole **μ** at an arbitrary azimuthal angle around the neurite, *ϕ*_neur_, can be computed by applying a rotation matrix to the components for a planar membrane:

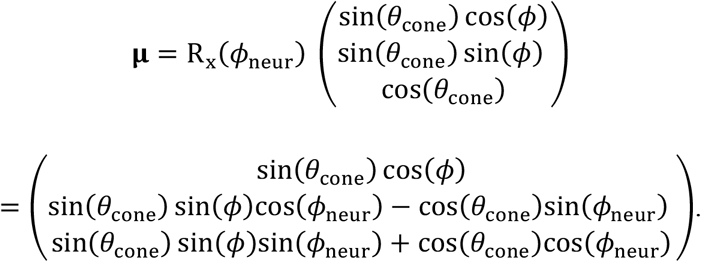

From this, one can evaluate the fluorescence excitation rate:

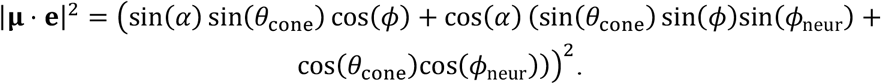

The fluorescence averaged over all chromophore orientations and around the neurite perimeter is:

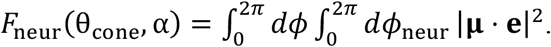

Evaluating the integral gives:

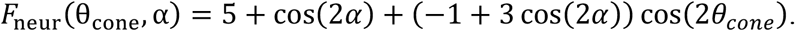

(An overall factor of 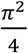 has been dropped because the overall scaling of the fluorescence is not important).

The FDLD signal depends on the fluorescence with polarization perpendicular to and parallel to the neurite axis. These are:

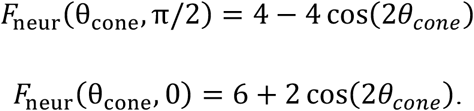

Inserting these expressions into the definition of FDLD gives:

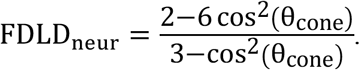

**Supplementary Figure 1.**
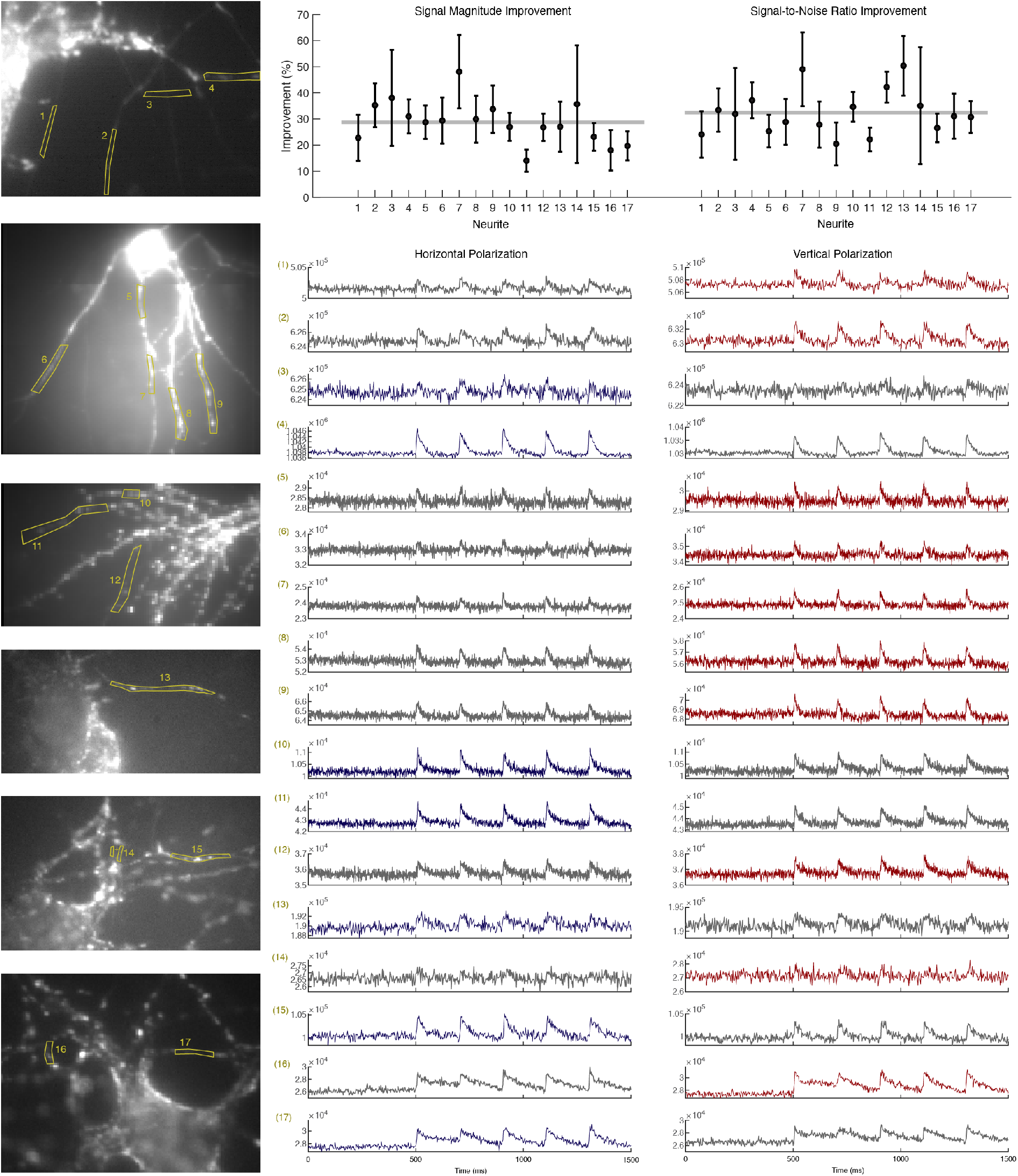
Examples of polarized excitation improving signal magnitude and SNR in neurites. Left: images of neurons with neurites selected which have predominantly vertical or horizontal orientation. Right: Fluorescence responses to induced action potentials taken under two polarizations. In traces for neurites illuminated with the preferred orientation are colored, traces with the orthogonal polarization are grey. Top: summary of data for each neurite.

